# An SNP variant MT1-MMP with a defect in its collagenolytic activity confers the fibrotic phenotype of Dupuytren’s Disease

**DOI:** 10.1101/2020.06.09.142513

**Authors:** Yoshifumi Itoh, Michael Ng, Akira Wiberg, Katsuaki Inoue, Narumi Hirata, Katiucia Batista Silva Paiva, Noriko Ito, Kim Dzobo, Nanami Sato, Valentina Gifford, Yasuyuki Fujita, Masaki Inada, Dominic Furniss

## Abstract

Dupuytren’s Disease (DD) is a common fibroproliferative disease of the palmar fascia. We previously identified a strong association with a non-synonymous variant (rs1042704, pD273N) in *MMP14* (encoding MT1-MMP). We investigated the functional consequences of this variant, and demonstrated that the variant MT1-MMP (MT1-N_273_) exhibits only 17% of cell surface collagenolytic activity compared to the ancestral enzyme (MT1-D_273_). Cells expressing both MT1-D_273_ and MT1-N_273_ in a 1:1 ratio, mimicking the heterozygous state, possess 38% of the collagenolytic activity compared to the cells expressing MT1-D_273_, suggesting that MT1-N_273_ acts in a dominant negative manner. Consistent with this hypothesis, patient-derived DD myofibroblasts expressing MT1-N_273_ demonstrated around 30% of full collagenolytic activity regardless of the heterozygous or homozygous state. 3D-molecular envelope modelling using small angle X-ray scattering demonstrated altered positioning of the catalytic domain within dimeric molecules. Taken together, our data suggest that rs1042704 directly contributes to the fibrotic phenotype of DD.

## Introduction

Dupuytren’s disease (DD [MIM 126900]) is the most common inherited disorder of connective tissues. It is a fibroproliferative disorder of the hand affecting approximately 4% of the general population, increasing to 20% in those over 65 years old in the UK, and presents to a wide variety of physicians and surgeons (Hart & Hooper, 2005; Lam *et al*, 2010). Progression of DD proceeds with the appearance of initial nodules followed by the formation of cord, which causes flexion contractures of the fingers, resulting in impairment of hand function. The cord is composed of myofibroblasts and fibrillar collagen (types I and III) (Lam *et al.*, 2010). Currently, the mainstay of treatment for DD is surgery (Misra *et al*, 2007), though other modalities are emerging (Hurst *et al*, 2009; Nanchahal *et al*, 2017). However, the mechanism of development of DD is still unclear, and even after adequate primary treatment, the recurrence of the disease is common (van Rijssen *et al*, 2012). In an attempt to understand the mechanism of DD, we have previously undertaken genome-wide association studies (GWAS) and found association at 26 independent variants (Dolmans *et al*, 2011; Ng *et al*, 2017). Candidate genes at these loci include those involved in WNT signaling, cell adhesion, matrix modulation, and inflammation (Dolmans *et al.*, 2011; Ng *et al.*, 2017).

Among these variants, rs1042704 was of particular interest, as it was the only exonic, protein-coding variant discovered in the GWAS. By using Bayesian analysis of 99% credible sets, it was concluded that this SNP is likely to be the causative variant at that locus (Ng *et al.*, 2017). rs1042704 is in the *MMP14* gene, which encodes the protein Membrane-Type 1 Matrix Metalloproteinase (MT1-MMP/MMP-14). The ancestral allele at rs1042704 is G encoding Asp_273_ (D_273_), while the alternate allele is A encoding Asn_273_ (N_273_) in the enzyme, where the A allele associates with DD. The allele frequency of the A allele in western European populations is around 0.20. To date, all reported functions and activities of MT1-MMP have been based on the common G allele gene product, with D_273_(Sato *et al*, 1994). MT1-MMP is a type I transmembrane proteinase that belongs to matrix metalloproteinase (MMP) family of enzymes (Itoh, 2015). It degrades extracellular matrix (ECM) components directly on the cell surface including fibrillar collagens type I, II, III (Ohuchi *et al*, 1997). MT1-MMP activates other MMPs on the cell surface including proMMP-2 and proMMP-13, expanding the proteolytic repertoire, and it also cleaves cell adhesion molecules such as CD44, modulating cellular adhesion to the matrix (Itoh, 2015). MT1-MMP is a crucial cellular collagenase *in vivo*, and MT1-MMP null mice display skeletal developmental defects and general fibrosis in soft tissues (Holmbeck *et al*, 1999). It has also been shown that MT1-MMP is the only collagenolytic MMP that directly promotes cellular invasion into the collagenous matrix, for example in cancer metastasis (Holmbeck *et al*, 2003; Holmbeck *et al*, 2004; Hotary *et al*, 2002; Miller *et al*, 2009; Sabeh *et al*, 2004). MT1-MMP was previously shown to be overexpressed in DD nodules, and MT1-MMP knockdown in DD-derived myofibroblasts was shown to reduce the contractility of the cells (Wilkinson *et al*, 2012). However, the precise role of MT1-MMP in the development of DD is not understood.

In this study, we have characterized biochemical and structural feature of the MT1-MMP SNP rs1042704 and found that the enzyme with N_273_ possesses significantly lower collagenolytic activity compared to the enzyme with D_273_ while fully retaining proteolytic activity against gelatin. Myofibroblasts from DD patients with G/G genotype that express MT1-MMP with D_273_ can degrade collagen efficiently, while cells derived from heterozygous (G/A) and homozygous (A/A) variant genotypes that express MT1-MMP with N_273_ degrade collagen in a significantly less efficient manner. 3D molecular envelope model of the ancestral and variant enzymes showed that the substitution of D_273_ to N causes significant influences in the arrangement of the ectodomains. Since MT1-MMP is thought to play a major role in soft tissue collagenolysis, our data suggest that rs1042704 plays a pivotal role in the fibrotic phenotype of DD.

## RESULTS

In Figure 1A, crystal structure model of MT1-MMP catalytic domain is shown (PDB: 1BUV). D_273_ is within the C-terminal helix of the catalytic domain, located on the surface of the molecule, and is distant from the catalytic site. Therefore, the substitution of D_273_ to N was not predicted to impact on either catalytic domain folding or the catalytic activity of the enzyme molecule. The structural model between the D_273_ and N_273_ indicated that N occupies a similar molecular space and is likely to change only the charge on the molecular surface (Figure 1B).

**Figure 1.**
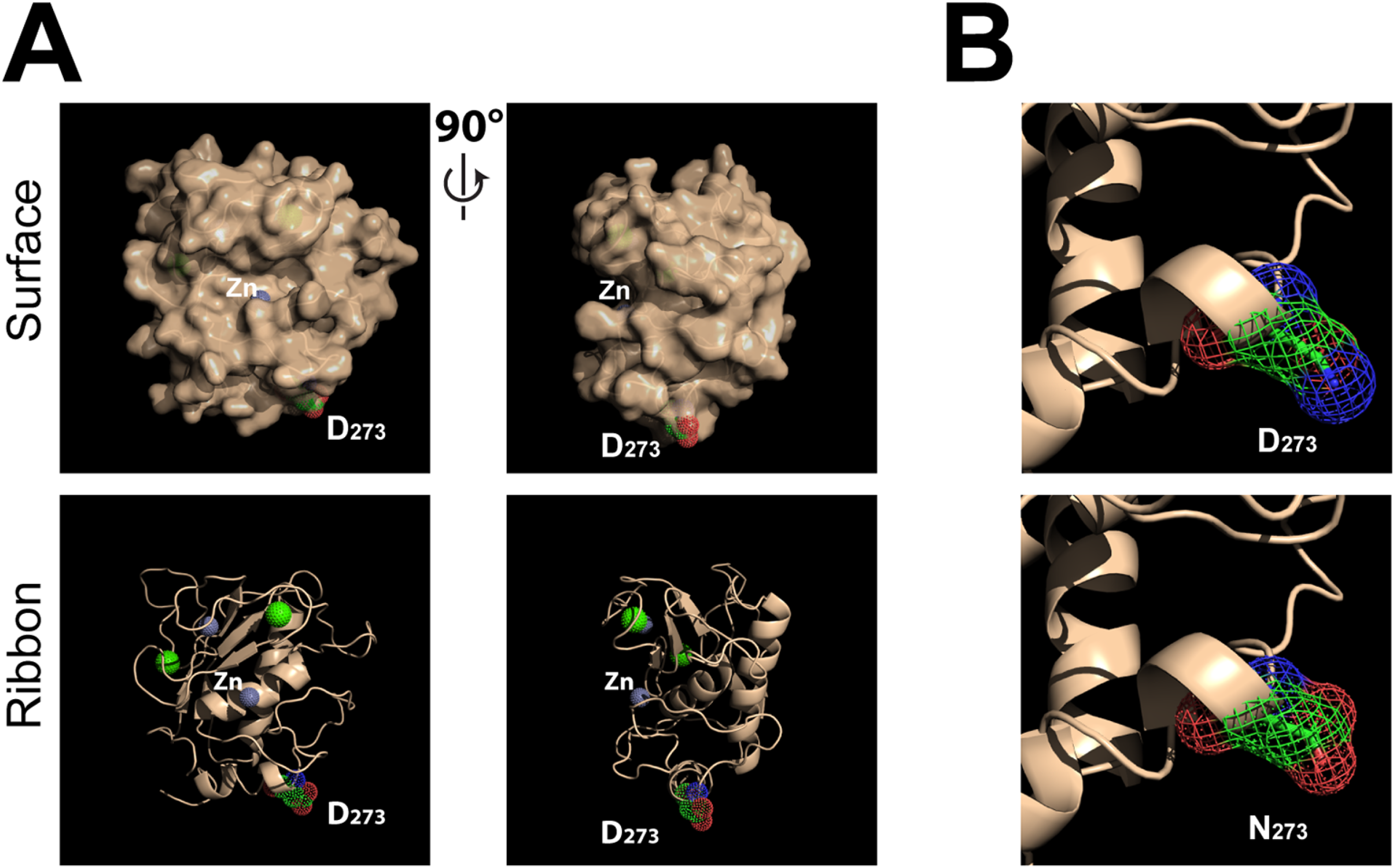
Position of Asp_273_ within the catalytic domain of MT1-MMP. **A.** MT1-MMP crystal structure (1BUV). Asp_273_ (D_273_), Zn, and Ca were highlighted as a sphere using PyMol. The structure is shown as a surface and ribbon model. The structure images on the right panels are rotated 90°clockwise. **B.** D_273_ was changed to N (Asn) in the PyMol software.

To investigate the properties of the SNP variant enzyme, we created the FLAG-tagged SNP variant construct (MT1-N_273_). First, a potential impact on general proteolytic activity was investigated. COS7 cells expressing FLAG-tagged MT1-MMP with D_273_ (MT1-D_273_) or MT1-N_273_ were subjected to a fluorescent gelatin film degradation assay. As shown in Fig 2A, cells expressing MT1-D_273_ or MT1-N_273_ degraded the gelatin film in an almost identical manner, suggesting that the SNP does not influence the catalytic activity of the enzyme, as predicted.

**Figure 2.**
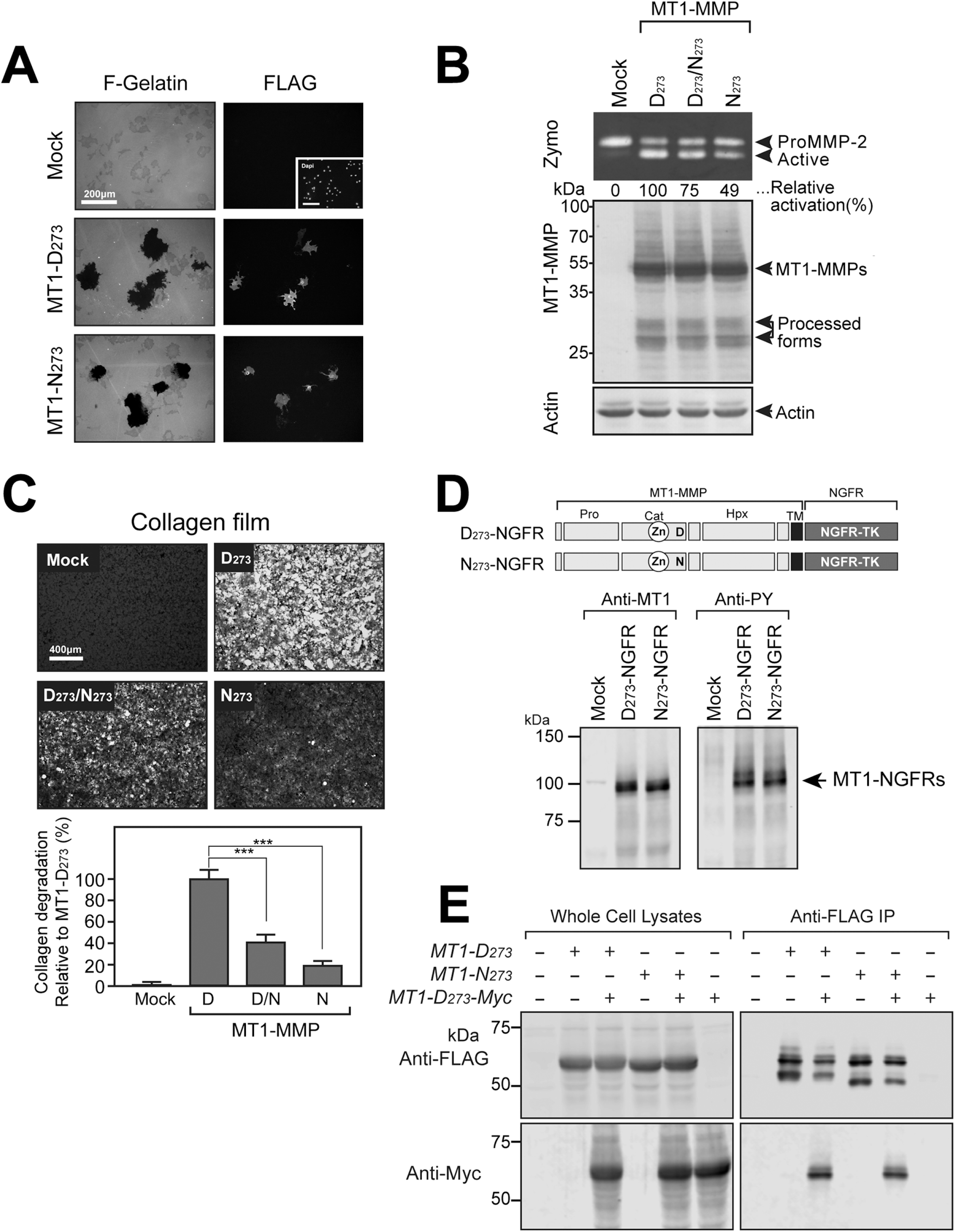
Effect of D_273_N substitution on biological activities of MT1-MMP. **A.** COS7 cells were transfected with expression plasmids for FLAG-tagged MT1-D_273_, MT1-N_273_ or empty plasmid, and subjected to fluorescently labeled gelatin film degradation assay. Cell surface MT1-MMPs were stained with anti-FLAG M2 antibody without permeabilizing cells. **B.** COS7 cells expressing MT1-D_273_ (D_273_) or MT1-N_273_ (N_273_) were cultured in proMMP-2-containing conditioned medium for 24h. ProMMP-2 in the media were analyzed by zymography (Zymo) and cell lysates by Western Blotting using anti-MT1-MMP Hpx domain antibody (MT1-MMP), and anti-actin as a loading control. **C.** COS7 cells expressing MT1-D_273_ or MT1-N_273_ were subjected to the collagen film degradation assay. Cells were transfected with MT1-D_273_ (D), MT1-N_273_ (N), both MT1-D_273_ and MT1-N_273_ in a 1:1 ratio (D/N), or empty plasmid (Mock). Images are representative of the collagen degradation activity of the cells, which is quantified in the bar chart. N=4 replicates for each transfection. Data represent mean ± SEM; ***p<0.01. **D.** COS7 cells were transfected with MT1-NGFR, and/or MT1-D_273_ and MT1-N_273_ as indicated and cultured for 48h. Cell lysates were subjected to Western Blotting for phosphotyrosine (PY) and MT1-MMP. Relative levels of PY were calculated by normalizing the intensities of PY bands with the intensity of the MT1-NGFR bands. **E.** COS7 cells were transfected with MT1-D_273_ and MT1-N_273_ together with or without Myc-tagged MT1-D_273_ (MT1-D_273_-Myc) as indicated. Cell lysates were subjected to immunoprecipitation with anti-FLAG IgG-conjugated beads. Whole cell lysates (left panel) and immunoprecipitated materials (right panel) were subjected to Western Blotting with anti-FLAG (top panel) and anti-Myc (bottom panel) antibodies.

We next examined the effect of MT1-MMP variants on proMMP-2 activation activity on the cell surface. MT1-MMP forms homodimer through both the hemopexin (Hpx) and the transmembrane domains (Itoh *et al*, 2006; Itoh *et al*, 2008; Itoh *et al*, 2001). ProMMP-2 activation depends on dimerization of MT1-MMP through the transmembrane domain (Itoh *et al.*, 2008), where One of the MT1-MMP molecules in the dimer forms a complex with TIMP-2 which acts as a receptor to attract proMMP-2 to the complex, while the other MT1-MMP cleaves proMMP-2 bound to the complex (Itoh, 2015). COS7 cells were transfected with MT1-D_273_ and/or MT1-N_273_, and cells were cultured in proMMP-2 containing medium for 18h. As shown in Figure 2B, MT1-N_273_ showed reduced activation of proMMP-2: 49% of the activation activity with MT1-D_273_. When the heterozygous genotype was mimicked by transfecting cells with a 1:1 mixture of MT1-D_273_ and MT1-N_273_ plasmids, the cells activated proMMP-2 with 75% activation efficiency (Figure 2B).

Next, we examined the collagen degradation activity. As shown in Figure 2C, COS7 cells expressing MT1-D_273_ efficiently degraded collagen film. In contrast, cells expressing MT1-N_273_ alone showed strikingly reduced collagen degradation: 17% of cells expressing MT1-D_273_. Interestingly when cells expressed both MT1-D_273_ and MT1-N_273_ in a 1:1 ratio, the collagen degradation was still significantly lower: 38% of the cells expressing MT1-D_273_ alone (Figure 2C). These data suggest that MT1-N_273_ may act in a dominant negative manner.

Cell surface collagenolytic activity of MT1-MMP depends on the enzyme to form a homodimer through the hemopexin (Hpx) domain (Itoh *et al.*, 2006). Therefore, we examined if MT1-N_273_ retains the ability to form a homodimer. We utilised the MT1-MMP constructs where its cytoplasmic domain was replaced with the cytoplasmic tyrosine kinase domain derived from nerve growth factor receptor (NGFR or Trk-A) (Figure 2D above, N_273_-NGFR and D_273_-NGFR, respectively). We have previously shown that when D_273_-NGFR forms a homodimer through either Hpx and/or transmembrane domains, the cytoplasmic domains of the D_273_-NGFR undergoes autophosphorylation (Itoh *et al.*, 2006; Itoh *et al.*, 2008; Itoh *et al.*, 2001). As shown in Figure 2D, expression of either D_273_-NGFR or N_273_-NGFR in cells caused efficient auto-phosphorylation detected by anti-Phospho-tyrosine antibody (Anti-PY), suggesting that substitution of D_273_ to N does not influence the ability of homodimer formation.

Since expression of both MT1-D_273_ and MT1-N_273_ resulted in lower than expected collagenolytic activity (Figure 2C), we hypothesized that MT1-D_273_ and MTI-N_273_ would form heterodimers, and that this dimer form is unable to degrade collagen efficiently compared to homodimers of MT1-D_273_. To test this, we next examined if MT1-N_273_ can form a heterodimer complex with MT1-D_273_ by co-immunoprecipitation. As shown in Figure 2E, FLAG-tagged MT1-D_273_ and MT1-N_273_ were co-expressed with Myc-tagged MT1-D_273_ (MT1-D_273_-Myc), followed by immunoprecipitation with anti-FLAG beads. The data indicate that both MT1-D_273_ and MT1-N_273_ efficiently co-immunoprecipitated MT1-D_273_-Myc, suggesting that MT1-D_273_ and MTI-N_273_ indeed form a heterodimer efficiently.

We next analyzed fibroblasts isolated from the affected cord tissue of DD patients of known genotype at rs1042704: G/G encoding MT1-D_273_, heterozygous G/A encoding both MT1-D_273_ and MT1-N_273_, or homozygous A/A encoding MT1-N_273_ only. Firstly, we tested their ability to activate proMMP-2. Each group of the cells was treated with or without collagen I (100 μg/ml) to induce proMMP-2 activation via DDR2 signal activation (Majkowska *et al*, 2017). Cells with G/G genotype efficiently activated proMMP-2 upon treatment with collagen (Figure 3A, 3B). However, when G/A or A/A genotype cells were treated with collagen, they showed a reduction in proMMP-2 activation. Upon combining quantification data in each genotype, overall proMMP-2 activation was reduced by around 50% in both G/A and A/A genotypes (Figure 3B).

**Figure 3.**
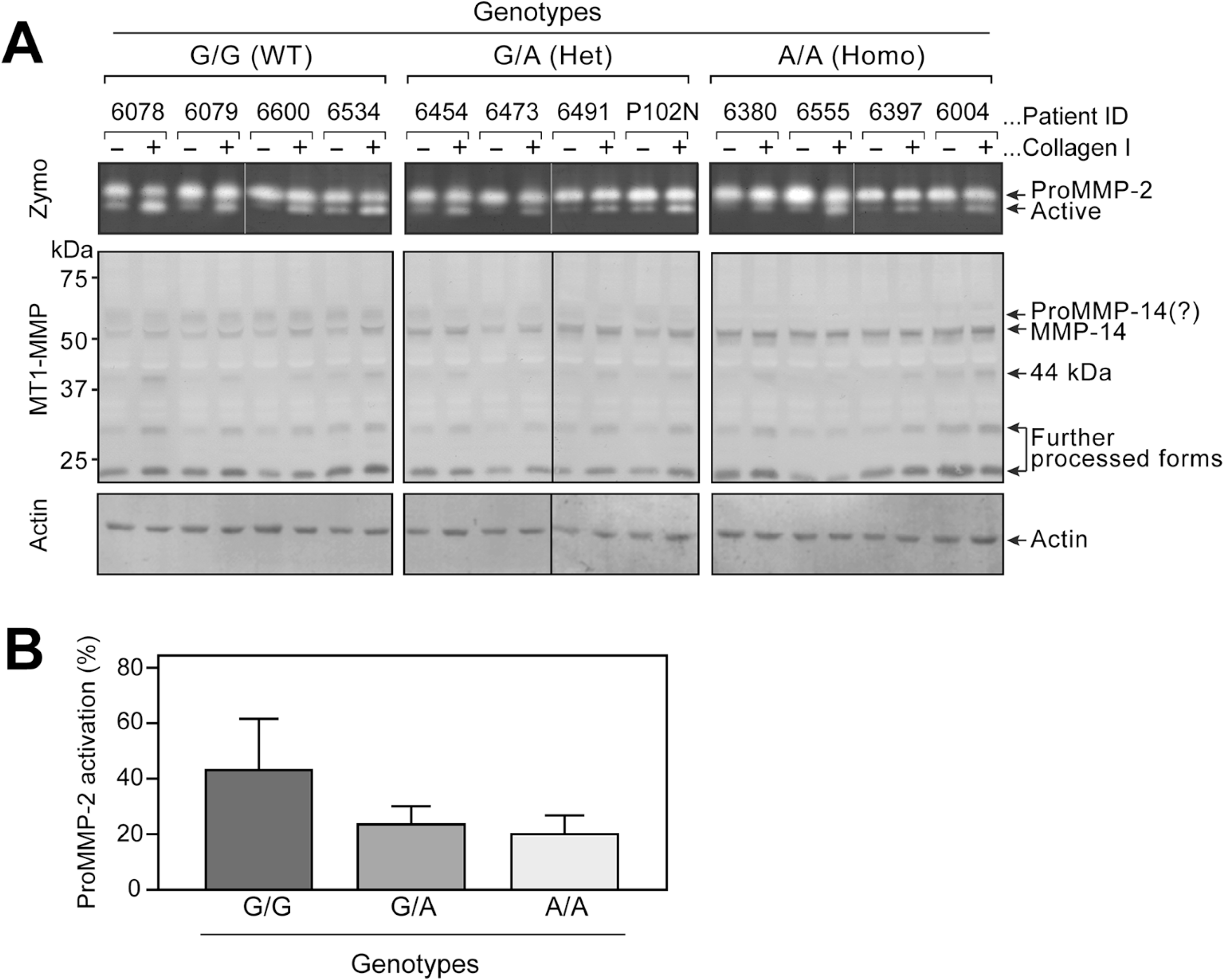
ProMMP-2 activation by different DD patient’s cells. **A.** ProMMP-2 activation by fibroblasts from DD patient tissues (4 different patient cells per genotype). Cells were cultured to the confluence in 12 well plates and stimulated with or without collagen I (100 μg/ml) in the serum-free medium for 24 h. ProMMP-2 and its active form were detected by subjecting the culture medium to zymography (Zymo). MT1-MMP was detected by Western Blotting using anti-MT1-MMP Hpx domain, compared to actin as a loading control. **B.** ProMMP-2 activation upon collagen stimulation of cells of each genotype group was quantified. N=4 replicates per genotype. The data represent mean ± SEM.

We next examined the cell surface collagenolytic activity of these cells. Patients’ cells with different genotypes were seeded on a thin film of type I collagen fibrils and cultured for 72h. All patients’ cells with G/G genotype effectively degraded the collagen film (Figure 4A). By contrast, all of the cells with G/A and A/A genotypes showed markedly reduced collagen degradation (Figure 4B). In aggregate, G/A genotype cells achieved 34% and A/A genotype cells 28% collagenolytic activity compared to G/G genotype cells (Figure 4C).

**Figure 4.**
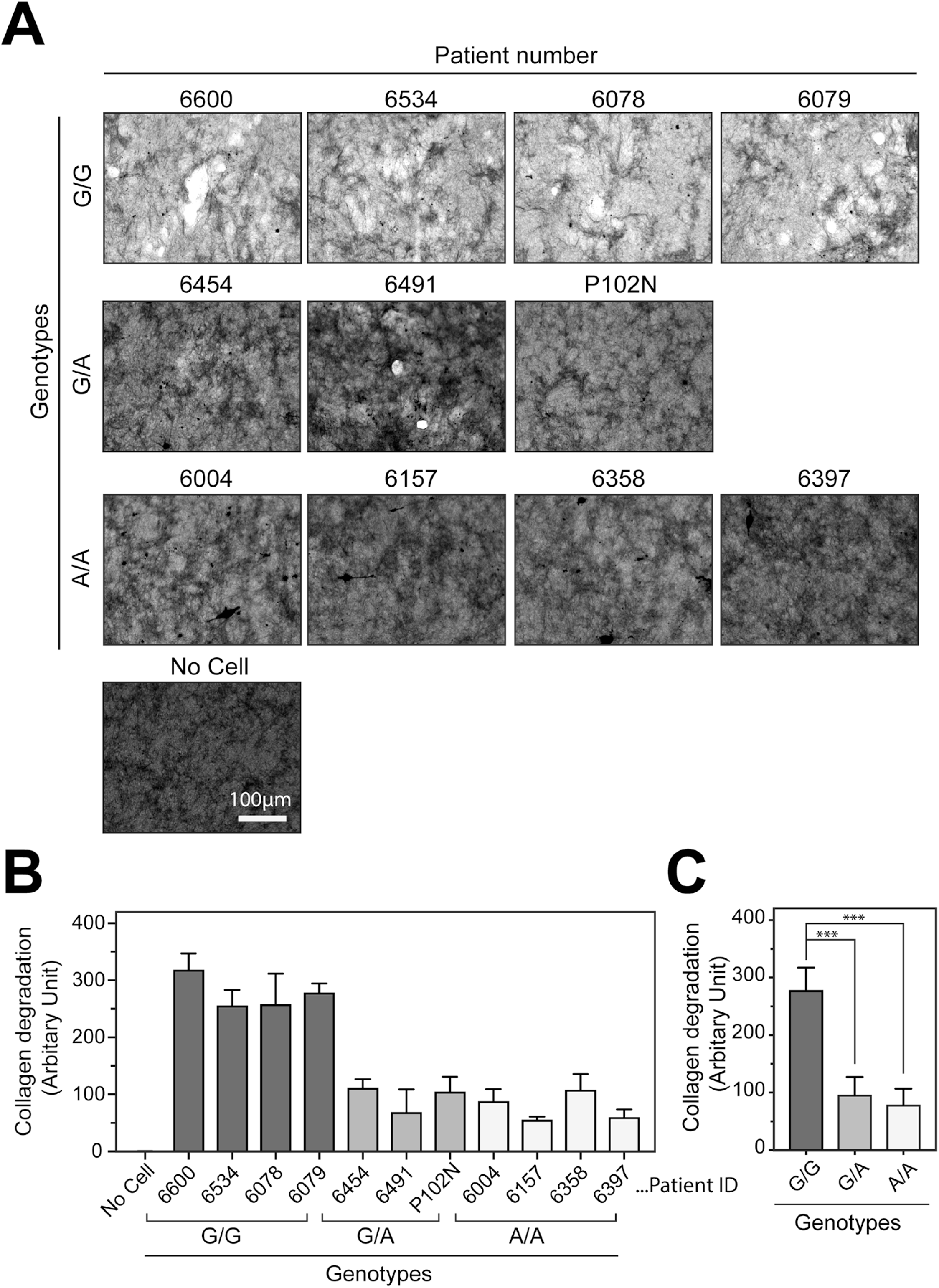
Collagen film degradation activity by DD patient’s cells. **A.** Patient’s cells were subjected to the collagen film degradation assay. Cells were cultured on collagen film in 12 well plates for 48h. A representative image from each patient’s cells is shown. **B.** Collagen degradation was quantified by image analyses, shown in the left panel as individual patient data (N=3 replicates for each patient’s cells). The data represent mean ± SEM. **C.** The graph demonstrates the aggregated data from panel B per genotype. The data represent mean ± SEM. ***p<0.01.

When we analyzed MT1-MMP in DD patient-derived myofibroblasts by Western blot (Figure 3), we noted that MT1-MMP of different genotypes displayed bands of slightly different molecular sizes (Figure 3A). G/G cells displayed a band at 62 kDa and a sharper band at 55 kDa. In A/A cells, the primary band was at 52 kDa, and the 62 kDa band was absent. In G/A cells, a mixture of molecular weight species from G/G and A/A cells was observed. The band intensities were also different, with the 52 kDa band in A/A genotype cells more intense than those at 62 and 55 kDa in G/G and G/A genotype cells.

We postulated that the different sizes and intensities were due to different processing of the enzyme molecule on the cell surface. MT1-MMP is known to be processed to lower molecular weight species due to auto-degradation, and metalloproteinase inhibitors such as GM6001 can inhibit this. Cells with G/G, G/A, and A/A genotypes were therefore cultured in the presence or absence of GM6001 for 24h and analyzed by Western Blotting. As shown in Figure 5A, neither the band pattern nor the intensities were affected by GM6001, suggesting that the different band patterns are not due to autolytic processing of MT1-MMP.

**Figure 5.**
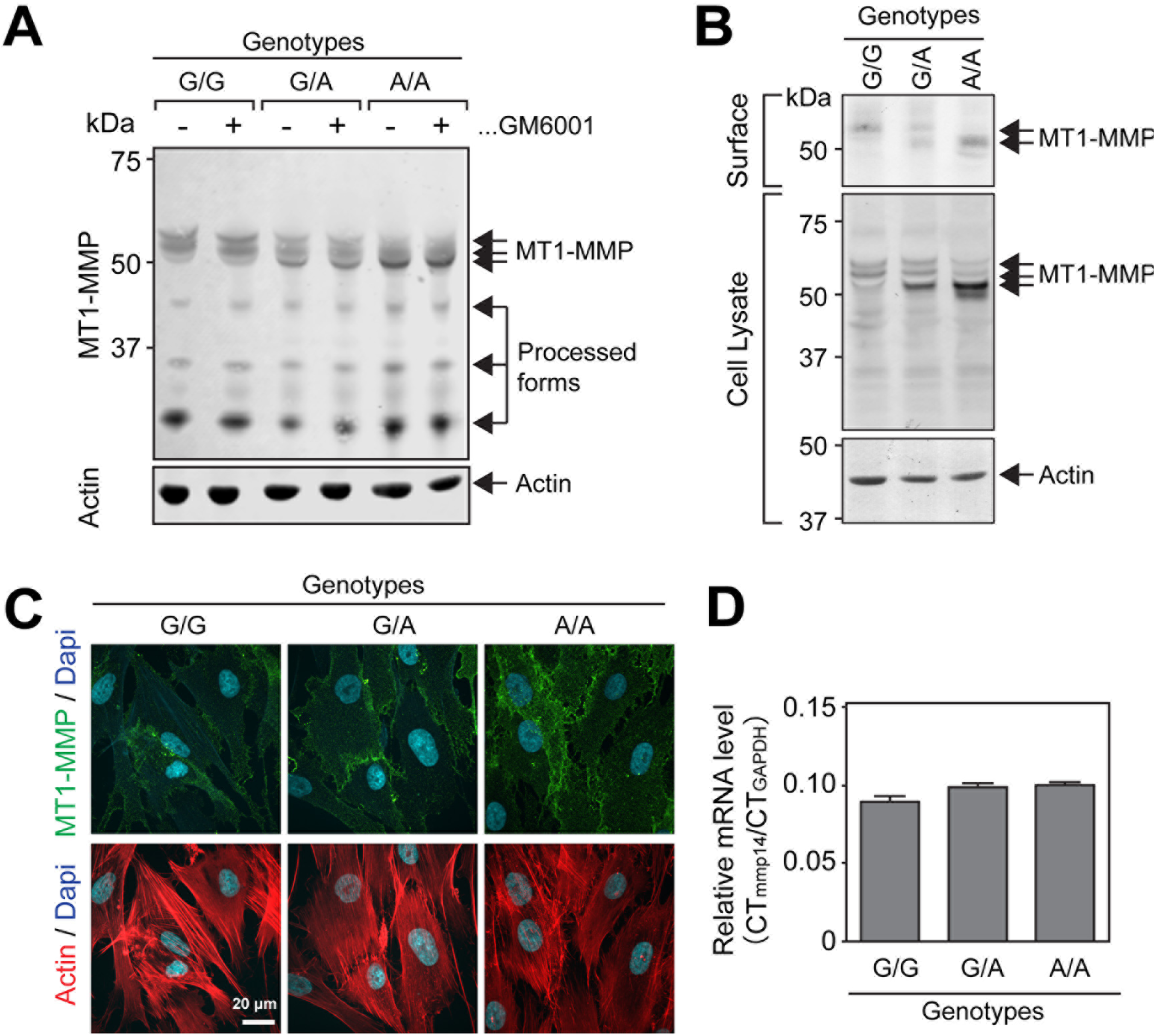
Analyses of endogenous MT1-MMP in patient’s cells. **A.** Cells from patient number 6078 (G/G), 6454 (G/A) and 6380 (A/A) were cultured to confluence in a 12 well plate in the presence or absence of GM6001 for 24h. Cell lysates were subjected to Western Blotting and stained for MT1-MMP and actin as a loading control. **B.** The same cells as in Figure 5A were subjected to cell surface biotinylation assay. **C.** The same cells were also seeded on gelatin-coated coverslips and subjected to indirect immunofluorescence staining without permeabilization. Fibrillar actin and nuclei were also stained. **D.** MT1-MMP mRNA levels in the same cells were analyzed by qPCR. The Y axis indicates the ratio of CT value obtained from qPCR for mmp14 and GAPDH (CT_mmp14_/CT_GAPDH_), thus, higher number inducates lower expression.

Next, we investigated if the level of band intensity reflected the cell surface level of the enzyme by subjecting cells to surface biotinylation (Figure 5B) and indirect immunofluorescence of cell surface MT1-MMP (Figure 5C). The data indicate that the cell surface level of MT1-MMP is notably higher in A/A genotype cells compared to G/G cells. We next examined if this was due to higher gene expression of MT1-MMP in the cells with A/A genotype by qPCR. As shown in Figure 5D, there were no significant differences detected in mRNA levels of MT1-MMP between the cells with different genotypes, suggesting that the difference in cell surface expression is not due to a difference in gene expression but instead may be attributed to post-translational events.

The position 273 is an N-terminal part of the C-terminal helix within the catalytic domain, and according to the structure of full length proMMP-1 (PDB:1su3) and the proMMP-2 (PDB: 1gxd), this part of the helix together with the hinge region forms an interface between the catalytic and the hemopexin domain. Therefore, substitution of D_273_ to N may influence the interface of these two domains and changes the orientation of the catalytic domain against the hemopexin domain, which would result in decreased collagenolytic activity. To test the hypothesis, we decided to analyze the 3D molecular envelope of the full ectodomain of MT1-MMP by small angle X-ray scattering (SAXS) method. To obtain a soluble form of MT1-D_273_ and MT1-N_273_ proteins, ectodomains of these enzymes were fused with the Fc portion of rabbit IgG, and expressed as soluble Fc fusion proteins (sMT1-D_273_-Fc and sMT1N_273_-Fc, Figure 6A). The Fc part of these chimera molecules enforces the ectodomain of the enzymes to form a disulfide bonds-mediated stable homodimer (Figure 6B), thus allowing the determination of the molecular shape of the MT1-MMP homodimer. These constructs were stably expressed in HEK293 cells in the presence of CT1746 to prevent auto-degradation, and the enzymes were purified from conditioned media using protein A agarose as described in the Methods. Purified sMT1-D_273_-Fc and sMT1N_273_-Fc were then subjected to size-exclusion chromatography SAXS (SEC-SAXS). We obtained a scattering curve on the elution peak from SEC, and the scattering curves from 2 variants were very similar (Figure 6C). The radius of gyration values (R_g *Guinier*_) were also consistent (Figure 6E). The molecular mass of each enzyme was estimated by calculating the volume-of-correlation from the scatting curve (Rambo & Tainer, 2013)(Figure 6E). The molecular mass of sMT1-D_273_-Fc was calculated to be around 140 kDa while calculated molecular weight from amino acids sequence was 123 kDa (Figure 6F upper panel). This difference is likely due to O-glycosylations present in the hinge (Wu *et al*, 2004) and relaxed conformational arrangement of each globular domains within the molecule. Interestingly the molecular mass of sMT1-N_273_-Fc calculated from SEC was 160 kDa, even bigger than the one from sMT1-D_273_-Fc (Figure 6F upper panel). We also calculated the pair distribution function, P(r) as shown in Figure 6D. The shape of P(r) functions were slightly different although Rg values (R_g *Pr*_, Fig 6E) were the same, suggesting that the overall shape of the molecules was similar but there is some differences in the conformation between the two molecules. Then, we have proceeded to the *ab initio* model calculation using GASBOR from the ATSAS package (https://www.embl-hamburg.de/biosaxs/software.html). Fourty individual models were obtained by separate calculations for each enzyme, and their mean positions were calculated and created the final molecular envelope models created (Figure 6G lower panels). The overall shapes of both sMT1-D_273_-Fc (blue) and sMT1-N_273_-Fc (yellow) are Y-shaped, where the elongated bottom region is likely to be the IgG Fc portion. However, there are notable differences in between them, especially in the top part of the molecule, which corresponds to the MT1-MMP ectodomain. Since both sMT1-D_273_-Fc and sMT1-N_273_-Fc have undergone processing of the pro-peptide upon secretion, the N-terminal part of the molecules is the catalytic domain (Figure 6B). Thus, it is clear that the position of the catalytic domains within the dimer form is not symmetrical. Since the final model is built from the average of 40 individual models, the region where they do not show a distinct domain shape or larger mass than the other is likely to reflect the non-rigid flexible positioning of the domain in the solution. Since it is not possible to register the catalytic or the hemopexin domain within the molecular envelope models due to their domain flexibility, it is still not clear how each domain is arranged. However, it is certain that change of the D_273_ to N has altered the flexibility and positioning of the catalytic and the hemopexin domains within the dimeric molecule, and this is likely the cause of significantly decreased collagenolytic activity of the MT1-N_273_ (Figure 2B).

**Figure 6.**
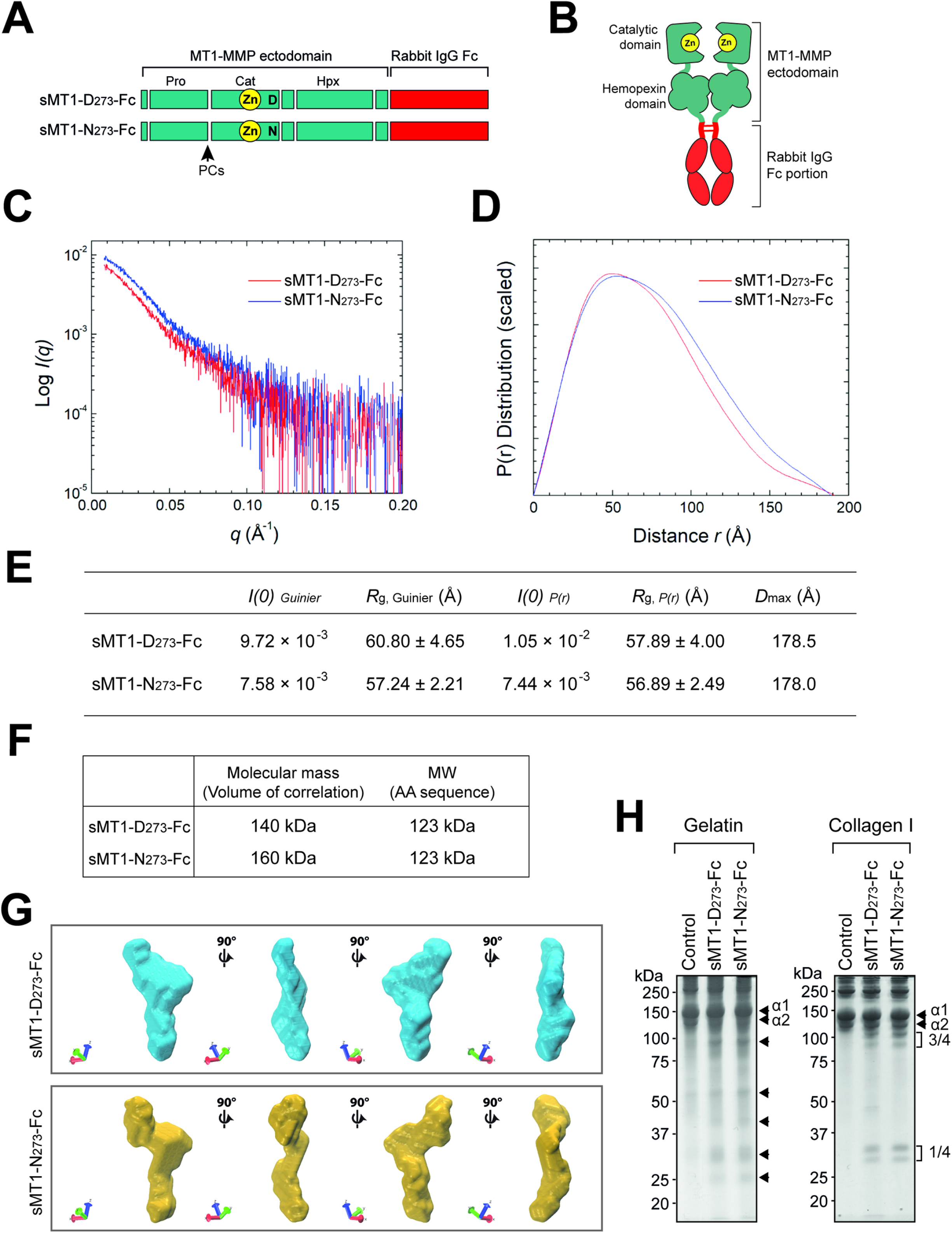
Analyses of 3D molecular envelope of soluble form of MT1-MMP variants. **A.** Schematic representation of the Fc-fusion MT1-MMP variants (sMT1-D273-Fc and sMT1-N273-Fc). Whole ectodomain of MT1-MMP variants (Met_1_-G_543_, green) were fused with rabbit IgG Fc portion (red). **B.** Schematic representation of sMT1-Fc protein. IgG Fc region mediates homodimer formation through the disulfide bonds. Pro-peptide is removed upon secretion by proprotein convertases (PCs), and the enzyme is secreted as active form. **C** Scattering curve of sMT1-D_273_-Fc and sMT1-N_273_-Fc (scattering intensity vs. scattering angle, q = 4πsinθ/λ). The SEC-separated sample was exposed to X-rays in a 1.6 mm diameter, 10 μm thick quartz capillary flow cell, followed by data collection every 3 seconds. Frames which were captured around the elution peak were selected, then the background subtraction and averaging were done. Scattering curves form 2 variants are similar. **D** Pair-distance distribution function (P(r)) for sMT1-D_273_-Fc and sMT1-N_273_-Fc. P(r) calculations were done by using ScÅtter. The overall shape of P(r) functions are slightly different between 2 variants although D_max_ values are almost same. **E.** Structural parameters which were derived from SAXS analysis **F.** The molecular mass calculated from the volume of correlation and amino acid composition for each Fc fusion MT1-MMP variant. **G.** The SEC-SAXS analyses of purified sMT1-D_273_-Fc and sMT1-N_273_-Fc. Top and lower panel shows the 90° rotated view of 3D molecular envelope models for sMT1-D_273_-Fc and sMT1-N_273_-Fc, respectively. The 3D envelope models were created by taking the mean of 40 individual models for each molecule. **H.** Analyses of gelatinolytic and collagenolytic activities of sMT1-D_273_-Fc and sMT1-N_273_-Fc. Purified sMT1-D_273_-Fc (17.5 ng) and sMT1-N_273_-Fc (17.5 ng) were reacted with gelatin for 4h at 37°C (left panel) or with type I collagen for 72h at 21°C (right panel). α1 and α2 chains for gelatin and collagen are indicated. Non-annotated arrows in the left panel indicate degraded fragments of gelatin. The ¾ and ¼ fragments of collagen α1 and α2 chains generated by collagenolysis are indicated for the right panel.

To address if the structural change is the direct cause of the reduced collagenolytic activity of the variant enzyme, we next compared gelatinolytic and collagenolytic activities of these purified enzymes. Since these enzymes were purified in the presence of CT1746, a hydroxamic acid-based broad spectrum MMP inhibitor, the purified enzyme fractions were dialysed extensively to remove CT1746, titrated against a known amount of TIMP-2, and subjected to the analyses. As shown in Figure 6H, incubation of these enzymes with gelatin in solution generated a similar degradation pattern and the levels of the degraded fragments, suggesting that change of D_273_ to N does not have an impact in the general proteolytic activity, which confirms the earlier data in Figure 2A. However, collagenolytic activity of sMT1-N_273_-Fc was almost identical to sMT1-D_273_-Fc as well, generating similar levels of ¾ and ¼ fragments of α1 and α2 chain of the collagen upon incubating at room temperature, which contrasts with the collagenolytic activity of MT1-N_273_ on the cell surface. The data suggested that the alteration of the structure does not influence their intrinsic collagenolytic activity, and the change of D_273_ to N causes the reduction of the collagenolytic activity only when the proteinase is expressed on the cell surface.

## Discussion

Fibrillar collagen type I is a major component of connective tissue, and is produced by fibroblasts. Under physiological conditions, the synthesis of collagen needs to be counterbalanced by its degradation, but when this balance tips towards more deposition of the collagen, which can be caused by more synthesis and/or less degradation of collagen, fibrotic conditions arise, such as is seen in DD. The proteolytic enzymes that are thought to be responsible for fibrillar collagen degradation include MMP-1, MMP-2, MMP-8, MMP-13 and MT1-MMP (Nagase *et al*, 2006). These enzymes can degrade fibrillar collagens at neutral pH. However, in animal models, only MT1-MMP knockout mice display a fibrotic phenotype of soft tissue (Holmbeck *et al.*, 1999), suggesting that MT1-MMP is the enzyme responsible for maintenance of collagen homeostasis.

Our data indicate that rs1042704 is associated with DD, and it was characterized as a likely causal variant (Ng *et al.*, 2017). Carrying the alternate allele increases the odds of a patient having DD by 1.25 and the allele frequency of 0.25 in cases means that approximately 43% of DD patients possess at least one copy of the alternate allele encoding the enzyme with N_273_. Among 26 loci associated with DD, rs1042704 was the only exonic SNP resulting in an amino acid substitution, D_273_ to N_273_ (Dolmans *et al.*, 2011; Ng *et al.*, 2017). Our data showed that this variant enzyme, MT1-N_273_, possesses significantly lower cell surface collagen-degrading activity, 17% of the activity of wild type. Interestingly, when MT1-D_273_ and MT1-N_273_ were expressed in a 1:1 ratio, cells possessed only 38% of collagen-degrading activity, suggesting that MT1-DN may act in a dominant negative fashion. We have previously shown that homodimerization of MT1-MMP is essential for its cell surface collagen-degrading activity through its hemopexin domain (Itoh *et al.*, 2006), and our data indicate that MT1-D_273_ and MT1-N_273_ can form heterodimeric complexes upon co-expression. Thus, it is possible that unexpectedly low collagenolytic activity upon co-expression may also be due to heterodimeric complex formation of MT1-N_273_ and MT1-D_273_ that has lower collagenolytic activity than a homodimeric complex of MT1-D_273_. When cells derived from patients with G/A genotype were analyzed, their collagenolytic activity was around 34% of G/G, almost as low as cells with A/A genotype (28%). While ectopically expressed MT1-N_273_ showed 17% of the collagenolytic activity of cells expressing MT1-D_273_, patients’ cells from A/A genotype exhibited 28% of collagenolytic activity. This may be attributed to higher cell surface level of the fully active form of the enzyme in A/A genotype cells compared to G/G genotype cells (Fig 5). Nevertheless, these data suggest that the impact of MT1-N_273_ on cellular collagenolytic activity is significant despite this higher cell surface expression, and regardless of it being present in the heterozygous or homozygous state.

The position of D_273_ is located in the N-terminal part of the C-terminal helix within the catalytic domain and is distant from the catalytic site. Therefore, it was unexpected that N_273_ negatively influences its collagenolytic and proMMP-2 activation activities. These two activities are not a simple catalytic function of the enzyme but require homodimerization of MT1-MMP on the cell surface which enables the enzyme to achieve specific molecular interactions (Itoh, 2015; Itoh *et al.*, 2006; Itoh *et al.*, 2001). Since the N-terminal part of this helix in the catalytic domain forms a part of the interface between the catalytic and the hemopexin domain according to the full-length structure of proMMP-1 and proMMP-2, we postulate that the D_273_ to N substitution would affect the orientation of the catalytic domain against the hemopexin domain.

The 3D molecular envelope models of sMT1-D_273_-Fc and sMT1-N_273_-Fc from SEC-SAXS analyses supported the hypothesis that the substitution of D_273_ to N influences orientation of the catalytic domain and the Hpx domain, which is likely the cause of decreased collagen degrading activity. However, there was no difference in intrinsic collagen degradation activity upon examination of the collagenolytic activity of these soluble enzymes in solution. The data suggest that differences are only have an effect when the enzyme is expressed on the cell surface. A similar finding was made when we previously examined soluble collagenase, MMP-13, previously (Itoh *et al.*, 2006). When MMP-13 was engineered to express on the cell surface by fusing MMP13 with the stalk region, the transmembrane and the cytoplasmic domains of MT1-MMP, this “membrane-type MMP-13” was completely inactive on the cell surface against the collagen film while it degraded gelatin (Itoh *et al.*, 2006). The membrane bound MMP-13 became an active collagenase only when an additional MT1-MMP hemopexin domain was further inserted to cause Hpx-domain-dimer(Itoh *et al.*, 2006). Thus, in the case of MT1-N_273_, although it is an active collagenase in solution as a soluble enzyme, the altered orientation of the catalytic and the Hpx domain somehow critically influences cell surface collagenolytic activity of the enzyme. It is possible that collagen degradation requires an additional molecular interaction on the cell surface, and the altered molecular conformation due to the D_273_ to N substitution might have caused a defect to this interaction. Further studies are necessary to fully reveal the mechanism.

The band pattern of ectopically expressed MT1-N_273_ on the Western blot is identical to MT1-D_273_ (Fig 2), however, when patient cells with G/G and A/A genotypes are compared, the band pattern of the endogenous MT1-N_273_ variant is different from MT1-D_273_. This is not due to variation between different individuals, but due to the genotype since the pattern is common among each genotype group. There are three major bands detected: 52 kDa, 58 kDa, and 62 kDa. In G/G genotype cells, the major bands are at 58 and 62 kDa, and the 52 kDa band was negligible. It is likely that the 58 kDa species is the active form. In A/A genotype cells the 52 kDa is the primary band, which is likely the active form. The band intensity of the A/A genotype cells is the highest of all regardless of the band species, and this reflects the amount of MT1-MMP at the cell surface in each genotype (Fig 5B & C). Since these different protein expression levels are not due to altered mRNA expression (Fig 5D), they are most likely caused by altered post-translational modification, attributable to the N_273_ substitution. The pattern of further processed forms of MT1-MMP that lack the catalytic domain are identical between different genotypes; thus, the difference is likely to be the catalytic domain or hinge region. It has been reported that MT1-MMP (D_273_) has four *O*-glycosylation sites at the hinge region (Wu *et al.*, 2004). The differences in molecular weight was also seen when MT1-D_273_ and MT1-N_273_ expressed in COS7 cells were immunoprecipitated (Figure 2E). Both pro and active form of MT1-N_273_ have lower molecular weight than MT1-D_273_. Therefore, it is possible that the substitution causes a different level of glycosylation at hinge region.

A thickened, contracted cord is the pathological feature that causes flexion contractures of the fingers and disability in DD. The cord is composed of myofibroblasts, and fibrillar collagens (types I and III) and the phenotype is considered to be due to the contractile nature of myofibroblasts and increased collagen deposition (Lam *et al.*, 2010; Townley *et al*, 2006). There are two possibilities that might cause excess collagen deposition: increased synthesis or decreased degradation, both of which would tip the balance of collagen metabolism and result in a fibrotic phenotype. As discussed above, complete knockdown of MT1-MMP in mice resulted in fibrosis of soft tissue (Holmbeck *et al.*, 1999), suggesting that defects in MT1-MMP-dependent collagen degradation are sufficient to result in fibrosis of soft tissues. Although to our knowledge collagen metabolism within cord tissue has not yet been investigated *in vivo*, we hypothesize that constant collagen turnover occurs in the cord tissue, and MT1-MMP plays an essential role in degradation. The strong association of rs1042704 and defects in collagenolytic activity of its gene product, MT1-N_273_, suggest that the phenotype of the myofibroblasts we characterized in this study may directly link to the thickening of the cord during the progression of DD.

According to the genome aggregation database (http://gnomad.broadinstitute.org), the allele frequency of rs1042704 in healthy general populations is approximately 0.2 in Europe and 0.16 worldwide, meaning that about 35% of Europeans and 30% of people worldwide carry at least one copy of the alternate allele. Since DD is a multifactorial polygenic disease (Ng *et al.*, 2017), carrying the alternate allele alone is not directly causative for DD, but it is expected to influence the ability of myofibroblasts in the palmar fascia to modify the cellular microenvironment significantly. This difference in allelic frequency between populations around the world might contribute to the different reported worldwide prevalence rates of DD (Hindocha *et al*, 2009).

In contrast to MT1-MMP null mice (Holmbeck *et al.*, 1999), people homozygous for the low activity allele (A/A) do not show any apparent developmental abnormality. This is likely because MT1-N_273_ retains some collagenolytic activity. Nevertheless, because this variant allele is present in a high proportion of the world’s population, further investigations aimed at understanding the mechanism of reduced collagenolytic activity is warranted. Furthermore, MT1-MMP represents an attractive therapeutic target in DD and related fibromatoses.

## MATERIALS AND METHODS

### Reagents

Mouse monoclonal anti-MT1-MMP Hpx domain (222-1D8) antibody was kindly provided by Prof Motoharu Seiki (University of Tokyo, Japan); rabbit monoclonal anti-MT1-MMP catalytic domain antibody (EP1264Y) was from Abcam (Cambridge, UK); actin antibody (C-19) was from Santa Cruz (Santa Cruz, CA, USA); anti-mouse and anti-goat alkaline phosphatase (AP)-conjugated antibodies were from Sigma-Aldrich (Dorset, UK); anti-rabbit AP-conjugated antibody was from Promega (Southampton, UK); Alexa 488-conjugated anti-mouse IgG and Alexa568-conjugated phalloidin were from Life Technologies (Paisley, UK); and GM6001 was from Elastin Products Company (Missouri, USA). A hydroxamic acid-based MMP inhibitor CT1746, and purified human TIMP-2 were kindly provided by Professor Gillian Murphy (University of Cambridge).

### Isolation of primary myofibroblasts

Primary myofibroblasts were isolated from surgical tissue of DD patients with fully informed consent. The study was approved by the Oxfordshire Research Ethics Committee: British Society for Surgery of the Hand Genetics of Dupuytren’s Disease (BSSH-GODD) study – B/09/H0605/65. Primary myofibroblasts were disaggregated from the fresh surgical tissues using 300 units/g type II collagenase at 1mg/ml (Worthington Chemical, NJ, USA) in Dulbecco’s Modified Eagle’s Medium (DMEM; Lonza, Slough, UK) supplemented with 5% fetal bovine serum (FBS; Labtech, East Sussex, UK) overnight at 37°C with 5% CO_2_. After incubation, cells were filtered using 40μm tissue culture strainer, frozen in DMEM containing 50% FBS and 10 % DMSO and kept until the experiments.

### Cell culture and transfection

COS7 cells were obtained from ATCC. COS7 cells were cultured in DMEM with 4.5g/l glucose supplemented with 10% DMEM and antibiotics (Lonza Biologics, Slough, UK). transiently transfected with expression plasmids using TransIT-2020 (Mirus, Madison, USA) according to the manufacturer’s instructions. Primary myofibroblasts were cultured in DMEM with 4.5g/l glucose supplemented with 10 % FBS and antibiotics.

### Plasmid construction

Flag-tagged MT1-MMP plasmid was constructed as described previously and subcloned into the pSG5 vector (Itoh *et al.*, 2001). FLAG-tag (DYKDDDDK) was inserted downstream of R_111_. This construct is named as MT1-D_273_ (or MT1-D_273_-F). Alternatively, Myc-tag (EQKLISEEDL) was inserted in the same position as described previously(Itoh *et al.*, 2001) (MT1-D_273_-Myc). The SNP variant MT1-MMP (MT1-N_273_) plasmid was constructed by mutating D_273_ to N within the MT1-D_273_ sequence using PCR-based mutagenesis and subcloned into pSG5 vector. The chimeric molecule of MT1-MMP and nerve growth factor receptor (Trk-A) was constructed as described previously (Itoh *et al.*, 2008). In this construct, the cytoplasmic domain of MT1-MMP (D_273_) was replaced with the one from NGFR containing the tyrosine kinase domain. The construct was subcloned into pSG5. Constructs to express soluble Fc fusion MT1-MMPs, sMT1-D_273_-Fc and sMT1-N_273_-Fc, were generated by inserting cDNA encoding the whole ectodomains of MT1-D_273_ or MT1N_273_ (cDNA encoding Met_1_-G_543_) into the multiple cloning site of pFUSE-rIgG-Fc1(InvivoGen). The cDNA fragment encoding sMT1-D_273_-Fc and sMT1-N_273_-Fc were then subcloned into pCEP4 vector (Thermo Fischer Scientific).

### Western blotting and zymography

Cell lysates were prepared in SDS-PAGE sample buffer (with 2-mercaptoethanol) and subjected to western blotting as described previously (Itoh *et al.*, 2006). Serum-free conditioned media were subjected to gelatin zymography (Itoh *et al.*, 2006). Densitometry analysis was performed in Image J software.

### Gelatin film degradation assay

Glass coverslips were coated with 50 μg/ml of Alexa Fluor 488-conjugated gelatin as described previously (Itoh *et al.*, 2006). Transfected COS7 cells (3×10^4^) were cultured on top of the coverslips for 6h. Cells were then fixed in 3% paraformaldehyde in PBS for 15 minutes and stained with DAPI (Sigma-Aldrich). Images were taken with Nikon Eclipse TE2000-E inverted fluorescence microscope equipped with a CCD-camera.

### ProMMP-2 activation assay

Cells were cultured to confluence in 12 well plates and stimulated with PureCol collagen (Advanced BioMatrix) at 100 μg/ml in the serum-free medium containing proMMP-2 for 24h. Culture media and cell lysates were harvested and analyzed by zymography and Western Blotting. ProMMP-2 activation was quantified by measuring the densities of the band of proMMP-2 and active MMP-2 using Image J and applying the data to the following equation: ProMMP-2 activation (%) = 100 × (density of active MMP-2) / [(density of proMMP-2) + (density of active MMP-2)].

### Collagen film degradation assay

Collagen film degradation was performed as previously described (Itoh *et al.*, 2006). Briefly, 12-well plates were coated with 100 μl of 2 mg/ml chilled and neutralized 1:1 mixture of PureCol collagen and Cellmatrix Type I-A collagen (Alpha Laboratories) and incubated at 37°C for 1 hour to induce fibril formation. COS7 cells were cultured on the collagen film for 24 hours and transfected with MT1-D_273_ and/or MT1-N_273_ plasmids in serum-free medium. Cells were further cultured for 48h, and attached cells were removed from the remaining collagen film by incubating in trypsin with EDTA in PBS (Sigma-Aldrich) extensively. The remaining collagen layer was fixed with 3% paraformaldehyde in PBS for 15 minutes, followed by staining with Coomassie Brilliant Blue-250. Images were taken with wide-field Nikon microscope operated by Volocity (Perkin-Elmer Life Sciences) with either a 10x or 4x objective lens. Quantification of collagen degradation activity was carried out by image analyses using Image J.

### Cell surface biotinylation assay

The cell surface biotinylation assay was carried out as described previously (Itoh *et al.*, 2001). Patient cells were cultured to confluency in a 100 mm dish, and cell surface proteins were labeled with sulfo-NHS-biotin (Thermo-Fischer), followed by affinity precipitation of biotinylated molecules by streptavidin Sepharose beads. The eluted samples were subjected to Western Blotting analyses using rabbit monoclonal anti-MT1-MMP antibody.

### Indirect immunofluorescent staining and imaging

Indirect immunofluorescence staining was carried out as described previously (Itoh *et al.*, 2001; Woskowicz *et al*, 2013). Briefly, myofibroblasts seeded on the gelatin-coated glass coverslips were fixed with 4% formalin in PBS for 5 min and blocked with 5% goat serum and 3% bovine serum albumin in PBS. Cells were then incubated with rabbit monoclonal anti-MT1-MMP antibody (Abcam, EP1264Y). Alexa Fluor 488- conjugated goat anti-rabbit antibody (Thermo-Fscher) was used to visualize the antigen signal. To visualize F-actin, cells were incubated with Alexa Fluor 568 phalloidin (Thermo-Fscher) in 0.1% Triton X-100 in TBS. Cell nuclei were visualized with DAPI. The fluorescent signals were imaged using Ultraview confocal microscopy (Perkin-Elmer Life Sciences).

### Co-immunoprecipitation

Co-immunoprecipitation was carried out as described previously (Itoh *et al.*, 2001). Briefly, COS7 cells were transfected with MT1-D_273_-F or MT1-N_273_-F with MT1-D_273_-Myc and cultured for 48h. Cells were then lysed in the RIPA buffer [1% Triton-X100, 0.1% SDS, 1% deoxycolic acid, 50 mM Tris-HCl(pH 7.5), 150 mM NaCl, 10 mM CaCl_2_, 0.02% NaN_3_] supplemented with proteinase inhibitor cocktail (Thermo-Fscher), and reacted with anti-FLAG IgG conjugated beads (Sigma-Aldrich) for 2h at RT. Bound materials were eluted by reacting the beads with 1 mg/ml FLAG peptide in TNC buffer [50 mM Tris-HCl (pH7.5), 150 mM NaCl, 10 mM CaCl_2_, 0.02% NaN_3_] at RT for 1h.

### Quantitative PCR (qPCR)

qPCR was carried out as described previously (Majkowska *et al.*, 2017). Total RNA was isolated from patient’s cells, and reverse-transcribed to cDNA with High-Capacity cDNA Reverse Transcription Kit (Life Technologies, Paisley, UK). MT1-MMP mRNA levels were measured by real-time quantitative PCR (qPCR) using the MT1-MMP TaqMan probe (Thermo-Fischer, Hs00237119_m1) and Ribosomal RNA Control Reagents (Thermo-Fischer). Relative changes in MT1-MMP mRNA were calculated using the comparative cycle threshold method.

### Expression and purification of sMT1-D_273_-Fc and sMT1-N_273_-Fc proteins

The pCEP4 plasmids with sMT1-D_273_-Fc and sMT1-N_273_-Fc were stably transfected into 293EBNA cells (Thermo-Fisher) and cultured in the growth media containing 400 μg/ml hygromycin and 10 μM CT1746 (provided by Prof Gillian Murphy)(Chander *et al*, 1995), a hydroxamate MMP-inhibitor, to prevent potential auto-degradation. To prepare conditioned media for purification of Fc fusion proteins, cells were cultured in the Corning® CellSTACK® Culture Chambers in the growth media with 400 μg/ml hygromycin. Upon confluency, cells were washed with PBS once and cultured in the serum free media containing 10 μM CT1746 without hygromycin for 7 days. The conditioned media were centrifuged to remove cell debris and subjected to protein A-agarose chromatography (Lifescience). After conditioned media were pass-through, the column was washed extensively with 50 mM Tris-HCl (pH7.5), 1M NaCl, 10 mM CaCl_2_, 0.02% NaN_3_ and Fc fusion protein bound to the column was eluted in 100 mM Glycine-HCl (pH3.0) which was immediately neutralised to pH8 by adding 2M Tris-HCl (pH8.0). All the fractions containing Fc fusion protein were corrected and dialysed against TNC buffer supplemented with 10 μM CT1746. Typically, 4 mg of purified Fc-fusion protein can be purified from 1L of conditioned medium.

### Size-exclusion chromatography small-angle X-ray scattering (SEC-SAXS)

The SEC-SAXS experiments were performed at beamline B21, Diamond Light Source (Didcot, UK), coupled with in-line size-exclusion chromatography. Protein samples were formulated at concentrations of 2.4 and 3.4 mg/mL for sMT1-D_273_-Fc and sMT1-N_273_-Fc respectively, in TNC buffer with 10 μM CT1746. Sample was loaded onto a Shodex KW404-4F column (4.6 mm ID × 300 mm) which was connected into Agilent 1200 HPLC system (Waters) at a flow rate of 0.16 mL/min. The SEC-separated sample was exposed to X-rays in a 1.6 mm diameter, 10 μm thick quartz capillary flow cell, followed by data collection every 3 seconds. It was confirmed on the size exclusion chromatogram system at B21 that the injected sample has been 3 times diluted at the exposure point. X-ray was focused on the detector, EIGER 4M (Dectris), the beam size was 1 mm (horizontal) × 0.5 mm (vertical) at the sample position and 0.08 mm (horizontal) × 0.07 mm (vertical) at the focal point. The wavelength of X-ray was 0.95 Å and the sample-detector distance was 2.7 m. The measurement temperature was 20 degrees centigrade. Raw SAXS 2-D images were processed with the DAWN (https://dawnsci.org/) processing pipeline at the beamline to produce normalised, integrated 1-D un-subtracted SAXS curves. The background subtraction, averaging of the data and determination of the structure parameters and the molecular mass were performed using the program ScÅtter (https://www.bioisis.net). On using ScÅtter, the molecular mass was derived from the volume of correlation which was directly calculated from the subtracted 1-D scattering curve (Rambo & Tainer, 2013). GASBOR from ATSAS software (https://www.embl-hamburg.de/biosaxs/software.html) was used for ab initio model calculation. 40 individual models were created for each species (sMT1-D_273_-Fc and sMT1-N_273_-Fc) and then those models were averaged with using DAMAVER package.

### Collagen and gelatin degradation in solution

Fractions containing purified sMT1-D_273_-Fc and sMT1-N_273_-Fc were dialysed against TNC buffer at 4°C to remove CT1746. The enzymes were titrated by known concentration of recombinant human TIMP-2 (provided by professor Gillian Murphy)(Crabbe *et al*, 1992) using a quenched fluorescent substrate: Mca-Pro-Leu-Gly-Leu-Dap (Dnp)-Ala-Arg-NH_2_, as described previously (Itoh *et al*, 1998). For gelatin degradation, neutralised PureCol collagen (2mg/ml) was denatured by incubating at 60 °C for 20 min. 20 μg portion of the gelatin was reacted with or without 17.5 ng of each enzyme at 37 °C for 4h in TNC buffer with 0.05% Brij35. After incubation, reaction was terminated by adding SDS loading buffer and boiled for 5min. Samples were then analyzed by 7.5% SDS-PAGE gel followed by staining with Coomassie Brilliant Blue R-250. For collagen degradation, PureCol collagen was neutralised (2mg/ml) and 20 μg portion of the collagen was reacted with or without 17.5 ng of each enzyme at room temperature (21°C) for 48h in TNC buffer with 0.05% Brij35. After incubation, reaction was terminated by adding SDS loading buffer and boiled for 5min. Samples were then analyzed by 7.5% SDS-PAGE gel followed by staining with Coomassie Brilliant Blue R-250.

### Statistical analyses

All quantitated data were plotted in GraphPad Prism software and the statistical significance was calculated using one-way ANOVA in the same software. At least 4 replicates for each treatment were incorporated into calculations. For proMMP-2 activation and collagen degradation, the mean ± SEM were shown.

## Acknowledgments

This work was supported by Kennedy Trust of Rheumatology Research Studentship (VG and YI), an Intermediate Clinical Fellowship from the Wellcome Trust (097152/Z/11/Z, DF), the NIHR Biomedical Research Centre, Oxford (DF), Japan Society for Promotion of Science Overseas Challenge Program for Young Researchers (NS), and Grant-in-Aid for Scientific Research (17H02157, MI), Long-term internship dispatch for innovation leader training (HN and MI), and Research Fellowship Abroad from São Paulo Research Foundation (2017/26813-9, KBSP). We thank Prof Gillian Murphy for providing CT1670 and human TIMP-2.

## Author Contributions

YI and DF conceptualize and oversaw the project. MN, AW carried out collection and genotypic characterization of the patients-derived myofibroblasts. KI carried out SEC-SAXS experiment and analyses. NH, KD, NS, VG characterized MT1-MMP activity of the patients-derived myofibroblasts. NI made constructs and characterised them. KBSP expressed and purify the MT1-MMP proteins. YS supervised NS. MI supervised NH.

